# A major chromosome 4 region modulates early vigor under chilling through brassinosteroid signaling associated genes in maize

**DOI:** 10.64898/2026.03.04.708938

**Authors:** Maxence James, Camille Clipet, Kristelle Lourgant, Benoît Decaux, Hélène Sellier-Richard, Delphine Madur, Sandra Negro, Stéphane Nicolas, Renaud Rincent, Alexandra Launay-Avon, Christine Paysant le Roux, Anca Lucau, Estelle Goulas, Andrea Rau, Catherine Giauffret

## Abstract

Early sowing is a key strategy to improve maize productivity and resilience under climate change, but it exposes plants to prolonged chilling stress that can severely compromise seedling establishment. While previous genetic studies have focused on germination or very early stages, tolerance to long-term chilling during the autotrophic transition remains poorly characterized. Here, we combined genome-wide association studies (GWAS) and transcriptome analysis on QTL near-isogenic lines (NILs) to dissect the genetic architecture of early vigor under chilling in maize. We identified a major genomic region on chromosome 4 (LD_COL4), harboring two QTLs within a 2.7 Mb interval, that were consistently associated with early vigor under long-term chilling conditions. Transcriptomic analysis of contrasted NILs revealed a cluster of differentially expressed genes co-localizing with LD_COL4, pointing to two strong candidate genes, *Zm00001d048582*, an ortholog of the Arabidopsis OPS gene that regulates the brassinosteroid (BR) signaling pathway upstream of the key transcription factors BES1 and BZR1, and *Zm00001d048612*, a brassinosteroid-signaling kinase (BSK). Multiple orthologs of BES1/BZR1 modulators were differentially expressed between genotypes under chilling, supporting the involvment of brassinosteroid signaling in this response. These findings highlight both genes as promising targets for marker-assisted breeding and gene editing to improve maize adaptation to early sowing.

## Introduction

Climate change and the reduction of agricultural inputs are destabilizing crop production. Maize, one of the most widely cultivated cereals in the world, is not immune to these new constraints, making the maintenance of its production and quality one of the most pressing challenges for global agriculture. Bringing sowing dates forward is one of the strategies being considered to meet this challenge, by optimizing solar and nitrogen availability in spring, managing the risk of heatwaves combined with increased water stress in summer, reducing fungal attacks in autumn, and limiting drying costs after harvest. However, maize exposed to temperatures below 10-15 °C can result in poor emergence rates, reduced seedling vigor, impaired photosynthetic efficiency, decreased leaf areas and ultimately substantial yield losses (Greaves, 1996; Leipner and Stamp, 2009). If breeding has substantially improved maize chilling tolerance to the extent that it can be cultivated at latitudes as high as 58° north, early sowing dates still require strong chilling tolerance, and in some oceanic regions like North Western Europe, tolerance to several weeks of chilling temperature is necessary. Moreover, the trend toward revisiting genetic resources to introduce new traits of interest (new end-uses, resistance to emerging pests, adaptation to intercropping practices, etc.) increases the risk of reintroducing cold-sensitivity alleles that had been eliminated from elite varieties. The ability to breed maize varieties adapted to these new agricultural challenges with enhanced chilling tolerance would improve overall biomass accumulation, thereby contributing to increased yields and improved feed and food security. Chilling tolerance in maize is a complex quantitative trait controlled by multiple genes, with different genetic mechanisms operating at distinct developmental stages and under varying environmental conditions. Plants respond to chilling stress through a sophisticated network of physiological and molecular adaptations, including the activation of cold-responsive genes, accumulation of soluble sugars, membrane lipid remodeling, metabolic adjustments, and the induction of reactive oxygen species (ROS) scavenging systems. Understanding the genetic architecture underlying these complex responses is essential for rapid and efficient breeding through marker-assisted selection and genomic approaches.

Over the past two decades, considerable progress has been made in identifying genetic loci associated with chilling tolerance in maize, initially through traditional QTL mapping approaches using bi-parental populations (Jompuk et al., 2005; Leipner et al., 2008) and more recently through genome-wide association studies (GWAS) based on large diversity panels (Strigens et al., 2013; Revilla et al., 2016). Very recently, the first genes involved in chilling tolerance have been deeply characterized, such as *COOL1,* a bHLH transcription factor that regulates downstream cold-responsive genes (Zeng et al., 2025) and *NLA*, a central regulator that links cold signaling to Pi homeostasis (Liao et al., 2026).

While genetic approaches such as GWAS provide powerful tools for identifying genetic variants associated with chilling tolerance, they offer limited insights into the dynamic molecular mechanisms and gene regulatory networks that underlie stress responses. Transcriptomic approaches, particularly RNA sequencing (RNA-seq), enable comprehensive characterization of genome-wide gene expression changes in response to environmental stresses, revealing temporal patterns of transcriptional reprogramming, the identification of stress-responsive genes, and the elucidation of regulatory pathways. Previous transcriptomic studies in maize have demonstrated that chilling stress triggers extensive transcriptional changes affecting thousands of genes involved in diverse biological processes, including signal transduction, hormone metabolism, carbohydrate metabolism, lipid biosynthesis, photosynthesis, antioxidant defense, and transcriptional regulation (Sobkowiak et al., 2014; Waititu at al., 2021).

Most genetic and genomics studies have been conducted under controlled conditions at the very early stages, from germination to emergence and rarely after the two-leaf stage. However, plants from susceptible genotypes can die at later stages (around the three- to four-leaf stage), if temperatures remain below 13°C after seed reserves are exhausted. In the field, this may compromise stand establishment even after fast and successful emergence. We used genome-wide association mapping to study the genetic architecture of the ability of plants to transition to autotrophy under long-term chilling conditions and a transcriptome analysis on QTL-near isogenic lines to generate hypotheses on the underlying genes and pathways. By integrating these complementary datasets, we aimed to identify high-confidence candidate genes The candidate genes identified in this study represent promising targets for marker-assisted breeding and gene editing.

## Materials and methods

### Genome wide association study

The analysis pipeline is summarized in Figure S1.

#### Plant material and genotyping data

This study was based on a Dent maize diversity panel assembled to analyze genetic groups relevant for hybrid breeding in Northern Europe, as described by Rincent et al. (2012). The panel consisted of 293 inbred lines genotyped using the Illumina MaizeSNP50 BeadChip array (Illumina Inc., San Diego, CA, USA), as described by Ganal et al., 2011. Quality control was performed to discard markers that did not meet the following criteria: less than 20% missing data, a minor allele frequency (MAF) greater than 1%, and heterozygosity lower than 15%. In addition, genotypes exhibiting more than 10% heterozygous calls were removed. The inbred lines were further genotyped using genotyping-by-sequencing (GBS) at Cornell University. GBS and imputation of missing data were performed as described by Elshire et al., 2011 and Swarts et al., 2014. Only biallelic markers were retained, and heterozygous calls were replaced with missing data. A second quality control step removed markers with a MAF lower than 1% and markers that became monomorphic after heterozygote removal. The MaizeSNP50 and GBS datasets were merged. For SNPs genotyped using both methods, the 50K genotypes were retained due to their higher accuracy and completeness. Among the 519,141 SNPs, GBS-derived markers with more than 70% missing data prior to Beagle imputation were excluded, based on comparisons of imputation results for approximately 7,000 SNPs common to both platforms. The final dataset was imputed using Beagle v4.1 (Browning and Browning, 2016) and consisted of 519,141 SNPs for 293 genotypes (Table S1).

#### Phenotyping data

Phenotyping was conducted on 293 hybrids obtained by crossing each Dent inbred line with UH007, a tester from the complementary Flint heterotic group. The panel was evaluated in three greenhouses during the winter season, with temperatures set to 14°C during the day and 9°C at night. Uniform emergence was achieved approximately one week after sowing by maintaining plants at 20 °C for the first four days, followed by thinning from three to two plants per pot at emergence. Each greenhouse contained approximately two-thirds of the hybrids in at least one replicate (one pot with two plants), while one-third of the hybrids was replicated once more within a greenhouse to enable spatial analysis. Replicated hybrids were distributed across three sub-blocks positioned along the expected temperature gradient within each greenhouse. Spatial heterogeneity was accounted for using a mixed linear model including genotype and greenhouse as fixed effects and sub-block nested within greenhouse as a random effect. The model was defined as follows:

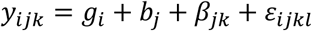

Where *y_ijk_* is the pot value for trait y, *g_i_* is the effect of genotype i, *b_j_* is the effect of greenhouse j, β_*jk*_ is the effect of sub-block k within the greenhouse j and ε_*ijkl*_ is the residual.

A similar model, with genotype treated as a random effect, was used to estimate variance components and broad-sense heritability. Heritability was calculated on an entry-mean basis as follows:

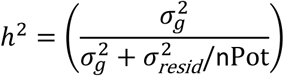

where 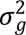 and 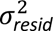 correspond to the genotypic and residual variance, respectively.

Several traits were recorded on the six to eight plants of each genotype. Early vigor (EV) was assessed by a visual scale from 0 for wilted plants to 5 for strong green plants, three (w3) and four (w4) weeks after emergence. The maximum length (LL) and width (WL) were measured on leaf #3, #4 and #5 after full expansion. The number of visible leaves (VL) and the shoot dry weight in g per plant (SDW) were evaluated at the end of the experiment, seven (w7) to eight (w8) weeks after emergence.

#### Genome-wide analysis

To reduce the ascertainment bias reported by Ganal et al. (2011), diversity analyses were restricted to SNPs selected from comparisons among nested association mapping founder lines (PANZEA SNPs; Gore et al., 2009). A total of 30,688 SNPs were retained. Kinship coefficients were estimated following Astle and Balding (2009) as follows:

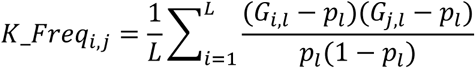

With *p_i_* the frequency of the allele 1 of the marker *l*, and *i,j* indicate the lines for which the kinship was estimated.

Population structure was inferred using ADMIXTURE v1.3 (Alexander et al., 2009), testing models with two to ten genetic groups. The optimal number of groups was determined based on principal coordinate analysis. Genome-wide association analyses were performed using a pipeline adapted from Negro (2019) calling FaST-LMM. Models accounted for relatedness using the kinship matrix (K) alone as it was sufficient to control false positives according to the QQ plots. To avoid loss of power due to strong linkage disequilibrium in centromeric regions, kinship matrices were computed using only markers located on chromosomes other than that of the tested SNP (Lisgarten et al. 2012; Rincent et al. 2014). Associations were tested between 516,630 SNPs and phenotypic traits with significance thresholds at a false discovery rate (FDR) of 5%.

Linkage disequilibrium (LD) was calculated as the squared allele frequency correlation (r²) between pairs of loci for each chromosome (Hill and Robertson, 1968). For the main target region, haplotype blocks were defined using Haploview (Barrett et al., 2005) with the confidence interval method of Gabriel et al. (2002). Only SNPs with MAF ≥ 0.05 were used for LD estimation. Genotypes were classified by haplotype to assess phenotypic differences among haplotype groups.

### Focus on the main target region for chilling tolerance

#### Construction, genotyping and phenotyping of QTL near-isogenic lines

Among ten available BC₅S₁ introgression populations derived from B73, the B73<CM174 population was selected because CM174 carried alternative alleles relative to B73 for all SNPs significantly associated with the major chilling tolerance QTL. For these SNPs, B73 carried the allele associated with improved early vigor despite its poor ability to transition to autotrophy under chilling conditions. Genotyping was performed using Kasp assays (LGC Genomics) for three markers within the target region. Five heterozygous BC2 plants for all or part of the region were identified and selfed. Subsequent genotyping with six additional markers guided the selection of candidate pairs. Final genotyping using a 25K SNP array identified two divergent S₁ individuals differing by a single introgressed segment spanning the entire target region. These lines were named M1 (B73 allele in the target region) and M2 (CM174 allele in the target region) and were selfed for phenotyping under chilling conditions.

The effect of the introgressed segment under chilling conditions was evaluated in three experiments (Exp1, Exp2, and Exp3). Seeds were germinated in Petri dishes at 20 °C in darkness for four days, then transplanted into pots filled with Klasmann TS3 substrate and transferred to a growth chamber or greenhouse for the chilling treatment. Exp1 was conducted in a growth chamber (day at 15.5 °C and 500 µmol photons/m^-^²/s^-1^ during 16 h; night at 10.5 °C) using ten plants per genotype and resulted in mortality for some individuals of the M2 line. Exp2 was conducted in a greenhouse during winter (day at 15°C during 16 hours, night at 11°C) using twelve plants per genotype, with milder chilling symptoms due to lower radiation. Exp3 was conducted in a growth chamber (day at 16°C and 500 µmoles photons/m²/s during 16h, night at 14°C) to avoid plant mortality and enable RNA-seq analysis, using 40 plants per genotype and replicate. Replicates corresponded to successive experiments conducted under identical conditions.

Several traits were evaluated starting from 20 days (d20) and up to 46 days (d46) after emergence. Early vigor (EV) was assessed by a visual scale from 0 for wilted plants to 5 for strong green plants. The number of visible leaf collars or leaf tips counted as well as the number of senesced leaf. The operating quantum yield efficiency of PSII (ΦPSII) was measured at a photon flux density of 500 µmol m^-2^ s^-1^ on the middle part of the youngest ligulated leaf using the miniaturized pulse-amplitude modulated fluorometer (MINI-PAM, Heinz Walz GmbH, Effeltrich, Germany). The shoot dry weight (SDW) was evaluated after drying at 70 °C for 72 h.

#### RNA-seq analysis

Exp4 under control conditions followed the same protocol as Exp3, but with temperatures set to 25°C during the day and 22°C during the night. Plants from both Exp3 and Exp4 were harvested at the three-leaf stage, and the third leaf was collected and split along the midrib. The first half was pooled for RNA-seq and the second half for further analysis. Twelve pooled samples (minimum of 30 plants per sample), corresponding to three biological replicates for each genotype and treatment, were collected and stored at −80 °C.

Total RNA was extracted from 100 mg of tissue using the NucleoSpin® RNA Plant kit and purified using the RNA Clean & Concentrator-5 kit. Libraries were prepared using the TruSeq Stranded mRNA protocol and sequenced on an Illumina NextSeq 500 platform using paired-end sequencing (75 bp reads), yielding approximately 15 million read pairs per sample.

Reads were trimmed using Trimmomatic v0.36 (Phred score > 20, read length > 30 bp), ribosomal RNA was removed using SortMeRNA (v2.1), and alignment was performed against the maize B73 version 4 reference genome using STAR (v2.5.2b). Gene expression was quantified using uniquely mapped paired-end reads, with multi-mapped reads discarded. On average, 89.77% of read pairs were assigned to genes. All steps of the experiment, from growth conditions to bioinformatic analyses, were uploaded to the CATdb database (Gagnot S. et al. 2007, http://tools.ips2.u-psud.fr/CATdb/).

Differential analyses were performed with a negative binomial generalized linear model including fixed effects for genotype, treatment, genotype × treatment, replicate, and a trimmed mean of M values (TMM) offset to normalize library sizes in the edgeR Bioconductor package (V3.28.0, McCarthy et al., 2012).

RNA-seq expression data were validated using RT–qPCR analysis with 1 µl of 100× diluted reverse transcription product, 500 nM of forward and reverse primers and 1× SYBR Green PCR Master Mix (Bio-Rad) in a real-time thermocycler (AriaMx Real-Time PCR System, Agilent). The three-step program comprised an activation step at 95°C for 3 min and 40 cycles of a denaturing step at 95°C for 15 s followed by an annealing and extension step at 60°C for 40 s. Q-PCR amplifications were performed using specific primers for each housekeeping genes and target genes. For each sample, subsequent RT–qPCRs were performed in triplicate. The expression of the target gene in each sample was compared with the genotype M1 sample and calculated with the delta delta Ct (ΔΔCt) method using the following equation: relative expression = 2–ΔΔCt, with ΔΔCt = ΔCtsample – ΔCtcontrol and with ΔCt = Cttarget gene – Cthousekeeping gene (for calculations, the geometric mean was considered between the Ct of the housekeeping genes). Using this analysis method, relative expression of the target gene in the control sample (M1) was equal to one (Livak and Schmittgen 2001).

Uniprot accession number, protein sequences and Gene Ontology (GO) annotations were retrieved for all DEGs with the uniprot database (https://www.uniprot.org/). The Arabidopsis orthologue was done with Biomart Ensembl Plants (Yate et al., 2022). Because two strong candidate genes were related to brassinosteroid (BR) signaling pathway, gene models associated with GO:1900457 (regulation of BR mediated signaling pathway) and GO:0009742 (BR mediated signaling pathway) were extracted.

Additionally, maize orthologs for Arabidposis (BR) signaling pathway genes listed by Nolan et al. (2020) were retrieved through Biomart Ensembl Plants or PLAZA_v4.5. Both lists were compared to DEG list and used for enrichment analysis.

## Results

### Major loci found for early vigor and leaf width under long-term chilling conditions

A highly diverse panel of 293 Dent inbred lines crossed with a Flint tester was used for GWAS. Based on 30,688 highly polymorphic PANZEA SNPs, the parental inbred lines were divided into four main genetic groups (Figure S2). The Lancaster group was the most represented, comprising 148 genotypes including Mo17 with admixture coefficients above 0.7. The Iodent group, represented by 20 genotypes including PH207, was clearly separated along the first principal coordinate. A third group including Stiff Stalk lines such as B73 and B14a comprised 19 genotypes and was distinguished along the second axis. A smaller group of 11 European lines, including D06, was also identified.

All ten traits evaluated under chilling conditions exhibited high heritability (≥ 0.7; Table 1), reflecting the robustness of the experimental design. Early vigor showed strong variability (Table S2), ranging from approximately 1 in the least vigorous genotypes to a mean of 4 after three weeks of chilling or 3 after four weeks. Leaf size was also highly variable, with some genotypes exhibiting leaves up to 50% smaller than the most vigorous lines. The number of visible leaves seven weeks after emergence was the most stable trait, although the latest genotypes lagged by up to three leaves. These differences resulted in large variation in shoot dry weight, with the least productive genotypes producing only 4% of the biomass of the most productive ones.

**Table 1.** Genetic parameters for traits evaluated on a dent diversity panel under chilling treatment in a greenhouse experiment: population mean, minimum and maximum, mean number of replicates, genetic and environmental variances, heritability on an entry-mean basis (h²).

| Trait† | Mean | Minimum | Maximum | nRep | Genetic variance | Environmental variance | $h^2$ |
| --- | --- | --- | --- | --- | --- | --- | --- |
| EV_w3 | 4.03 | 1.35 | 5.45 | 3.25 | 0.356 | 0.508 | 0.69 |
| EV_w4 | 3.35 | 1.25 | 4.79 | 3.15 | 0.179 | 0.266 | 0.68 |
| LL3 | 285 | 191 | 380 | 3.06 | 640 | 424 | 0.82 |
| WL3 | 16.4 | 11.8 | 22.0 | 3.07 | 3.18 | 1.62 | 0.86 |
| LL4 | 399 | 270 | 492 | 2.75 | 1246 | 964 | 0.78 |
| WL4 | 24.7 | 15.9 | 35.0 | 2.75 | 9.39 | 3.34 | 0.89 |
| LL5 | 515 | 362 | 669 | 2.20 | 1762 | 1638 | 0.70 |
| WL5 | 36.1 | 26.4 | 51.7 | 2.19 | 16.58 | 7.54 | 0.83 |
| VLN_w7 | 6.75 | 5.27 | 8.16 | 3.13 | 0.134 | 0.172 | 0.71 |
| SDW_w8 | 3.18 | 0.22 | 6.06 | 3.26 | 0.715 | 0.695 | 0.77 |

† Traits are the early vigor (EV) assessed by a visual scale from 0 for wilted plants to 5 for strong green plants, the maximum length (LLn) and width (WLn) of leaf n in mm, the number of visible leaves (VLN), the shoot dry weight in g per plant (SDW). These traits were evaluated after 3 weeks (w3) to 8 weeks (w8) after emergence.

GWAS identified 58 significant SNP-trait associations (Figure 1; Table S3). Forty-two of these were associated with early vigor and defined a major 2.7 Mb region on chromosome 4. Seventeen of the 28 SNPs in this region were located within eight gene models from *Zea mays* B73 RefGen_v4.39 (Table S3). The most significant SNP (−log₁₀P = 8.8) was located 1.6 kb upstream of *Zm00001d048612* (4:1.286.635-1.292.972) while the second most significant SNP (−log₁₀P = 8.6) was located within the coding sequence of *Zm00001d048582* (4:218.617-220.062). Secondary loci for early vigor were found on chromosomes 1, 3, 5 and 10, located within the coding sequences of *Zm00001d033004, Zm00001d044517*, *Zm00001d018038* and *Zm00001d023404* respectively. For all SNPs associated with early vigor, B73-like alleles were associated with improved performance (Table S3). Their frequency in the 2.7 Mb region on chromosome 4 ranged from 71 % (*Zm00001d048585*, *Zm00001d048600*) to 92% (*Zm00001d048603*) in the full panel (Table 2). In *Zm00001d048582* and *Zm00001d048612,* it was around 80% in the full panel but dropped to approximately 35% in the Stiff Stalk group. This group had a much lower frequency of favorable alleles than the others for all early vigor positional candidates on chromosome 1 and 4. For the chromosome 3, 5 and 10 unique significant SNP, the frequency of the B73-like alleles in the panel was between 70% (chromosome 5) and 97% (chromosome 10). The lowest frequency was observed in the Lancaster, European and Iodent group respectively.

**Fig.1:**
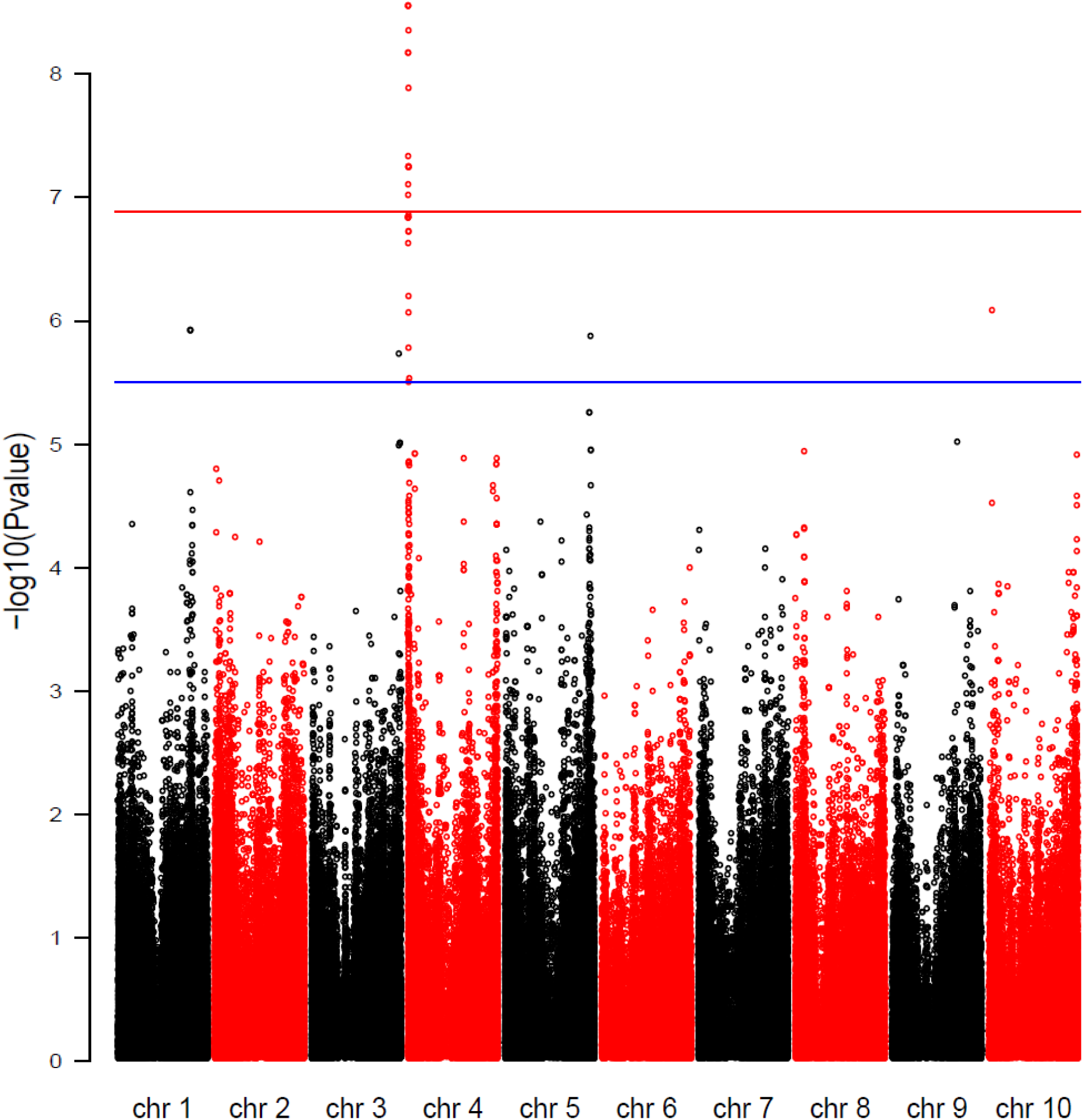
Manhattan plot for early vigor rating after 3 weeks of chilling treatment at 15°C during the day and 11°C during the night.

**Table 2.**
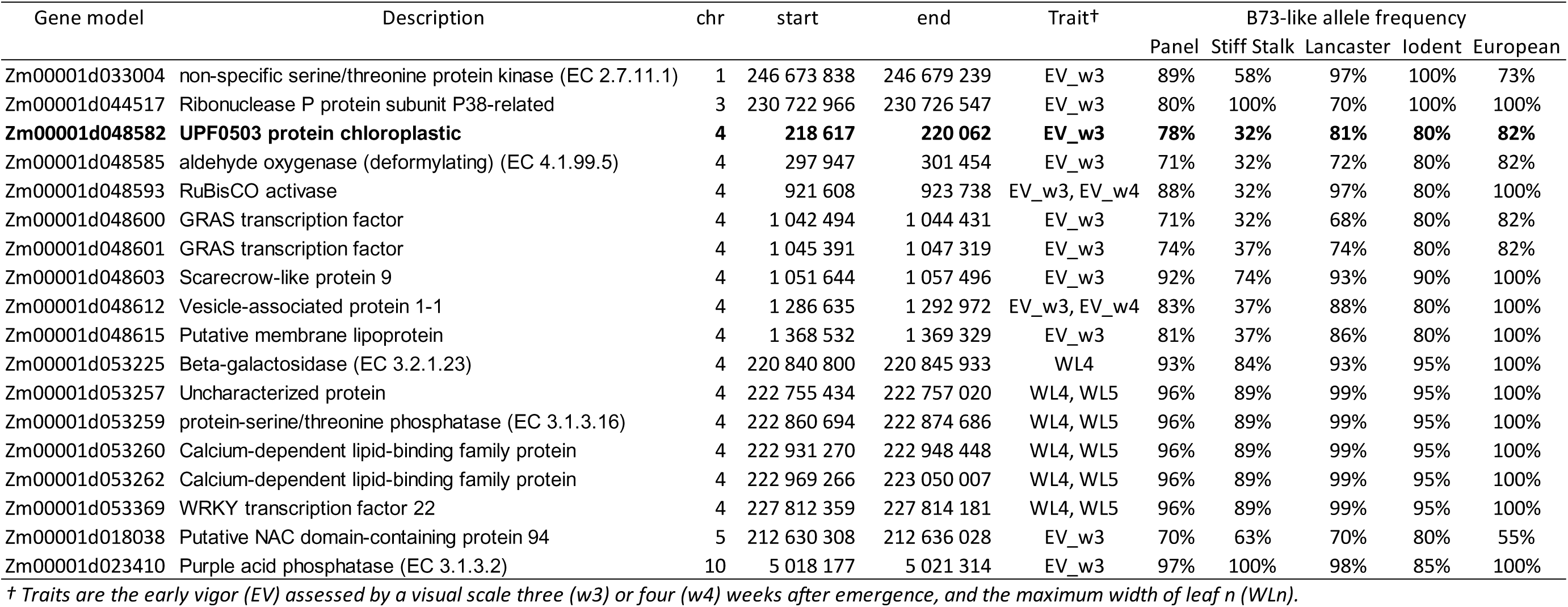
SNP-associated candidate genes for early vigor and leaf width and frequency of B73-like alleles in each genetic group. The panel comprised 293 dent inbred lines crossed to the UH007 flint tester. Each genetic group was represented by the subset of inbred lines with an admixture coefficient above 0.7.

A second region on chromosome 4 was associated with leaf width traits and spanned 7.0 Mb (Table S3; Table 2). *Zm00001d053257* and *Zm00001d053259* were the two genes associated with the peak of significance (−log₁₀P = 6.9). In contrast to early vigor, B73-like alleles in this region were associated with narrower leaves and were present in more than 90% of inbred lines.

### Two major QTLs for early vigor within the 2.7 Mb region on chromosome 4

Given the high level of significance and the large phenotypic variation observed, subsequent analyses focused on the 2.7 Mb region on chromosome 4, hereafter referred to as LD_COL4.

Linkage disequilibrium among significant SNPs in this region is shown in Figure 2A. Two distinct blocks of high LD were identified. The first block spanned positions 217,437 to 299,706 bp and was designated LD_COL4.1, while the second block extended from 1,285,003 to 1,369,532 bp and was designated LD_COL4.2. Of the 48 and 58 SNPs initially identified within LD_COL4.1 and LD_COL4.2, respectively, 35 and 42 SNPs were retained for haplotype definition (Table S4). For both regions, the B73-like haplotype was the most frequent among the female parents of the panel, represented by 94 individuals for LD_COL4.1A and 58 individuals for LD_COL4.2A. In the LD_COL4.1 region, seven additional haplotypes were shared by more than ten individuals. Four of these major haplotypes carried exclusively alleles associated with improved early vigor: LD_COL4.1B (W9-like), LD_COL4.1E (D06-like), LD_COL4.1F (H99-like), and LD_COL4.1H (Table S4). In contrast, two major haplotypes contained only alleles associated with reduced early vigor: LD_COL4.1C (B14a- or Mo17-like) and LD_COL4.1G (W117- or UH250-like). In the LD_COL4.2 region, five haplotypes, including the B73-like haplotype, were shared by at least ten individuals. Four of these haplotypes (LD_COL4.2B, LD_COL4.2D, LD_COL4.2E, and LD_COL4.2F) carried no alleles associated with reduced early vigor, whereas the LD_COL4.2C haplotype (Mo17-like) carried no alleles associated with improved performance (Table S4). Hybrids carrying haplotypes devoid of favorable alleles exhibited significantly poorer early vigor (Figure 2B, Figure S3).

**Fig.2:**
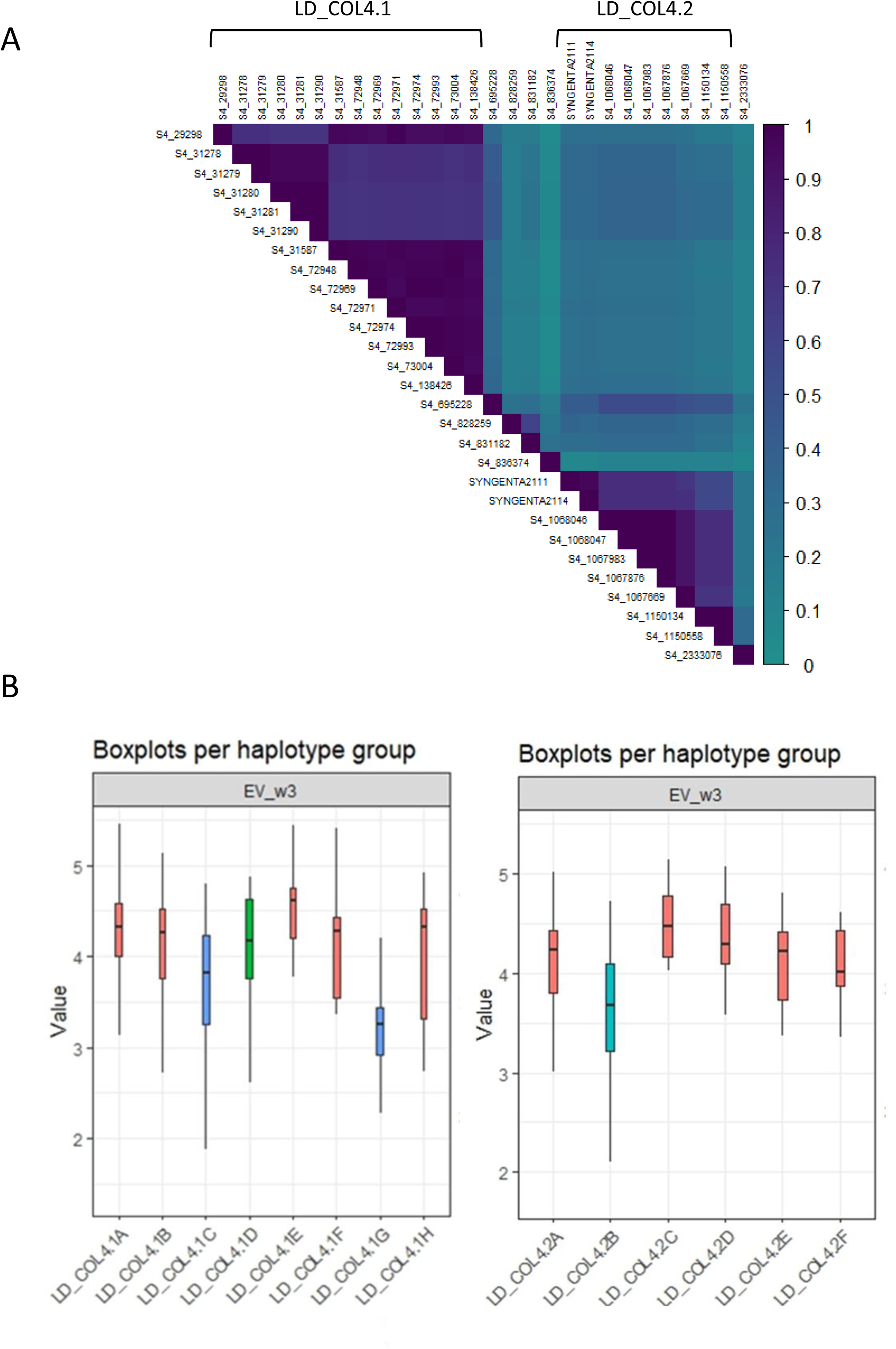
**A.** Linkage disequilibrium matrix for the LD_COL4 region linked to early vigor under chilling conditions. **B.** Effects of the LD_COL4 main haplotypes found in the diversity panel on early vigor.

### Phenotypic validation using QTL near-isogenic lines

Among the selfed progenies of five individuals from the BC5 population B73<CM174 showing residual heterozygosity within the LD_COL4 region, nine markers (Table S5) were used to select the pair of near-isogenic lines with the shortest CM174 introgression. Medium-density genotyping with the 25K SNP array demonstrated that the two lines differed by a single introgressed segment of approximately 5 Mb on chromosome 4. This unique segment spanned from PZE-104000008 (217,580 bp) to PZE-104006040 (5,146,805 bp), encompassing the entire 2.7 Mb LD_COL4 region. The contrasted phenotypes under long-term chilling conditions were confirmed in two growth chamber experiments and one greenhouse experiment. Depending on the stress intensity, the plants showed clear visible differences of chlorosis, wilting or senescence of the leaves (Figure 3). In addition to marked differences in early vigor at the one- to two-leaf stage (Table 3), significant genotypic differences were also detected for leaf developmental traits (number of visible leaf tips or collars, up to 29% reduction in M2), leaf senescence (up to 83% increase in M2), PSII photosynthetic efficiency, and shoot dry weight (23% reduction in M2 in Exp1).

**Fig.3:**
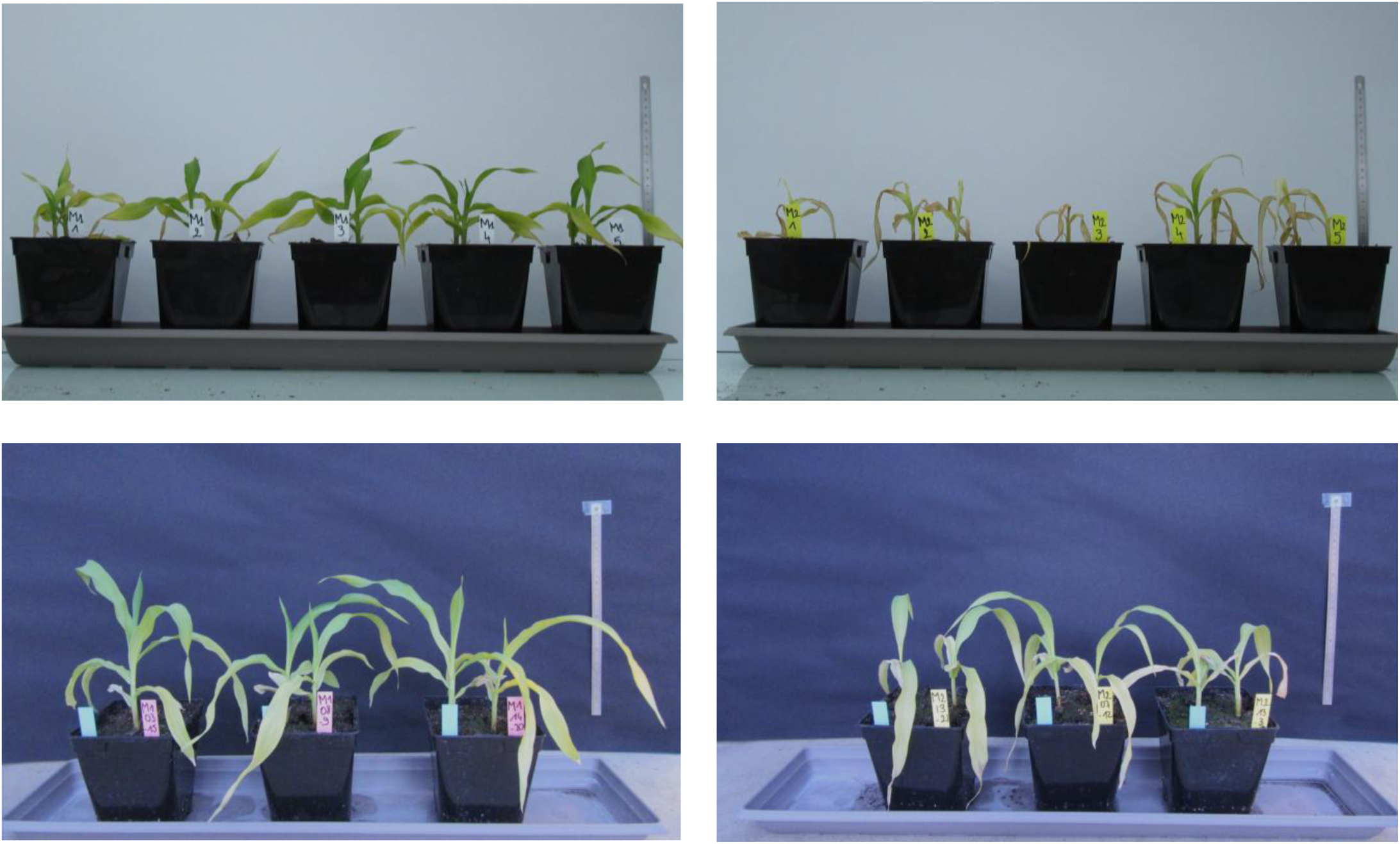
QTL-NILs for chr4 region after 28 days under chilling treatment in Exp1 (A, B) and Exp3 (C, D). M1-NIL (A, C) has the B73 allele, expected to confer a better vigor. M2-NIL (B, D) has the CM174 allele.

**Table 3.**
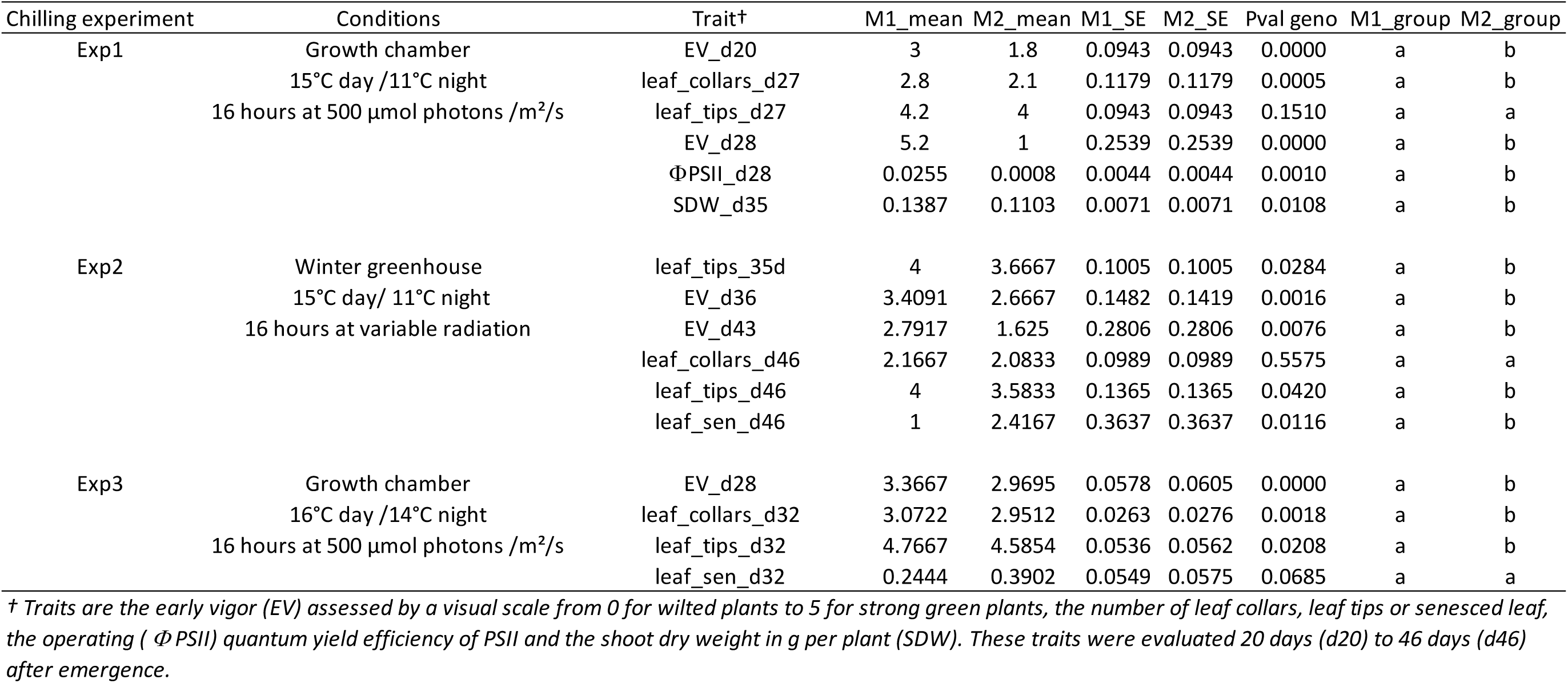
Phenotypes of two near-isogenic lines for a QTL related to chilling tolerance in 3 differents chilling experiments.

### Transcriptome analysis identifies a cluster of differentially expressed genes

The third chilling experiment was designed to induce clear phenotypic differences without causing plant or tissue mortality, thereby enabling sampling for transcriptomic analyses. This experiment was complemented by a control treatment at favorable temperatures. Under control conditions, no differences in plant vigor were detected between the two near-isogenic lines.

Mapping results were obtained for 44146 gene transcripts (Table S6) and 30322 of those were conserved after filtering for differential analysis. The transcripts of all 18 positional candidate genes identified by GWAS were successfully mapped; however, seven were excluded from further analysis due to low expression levels (Table S7).

Differential expression analyses were conducted to assess the genotype effect within each treatment and the treatment effect within each genotype, as illustrated in Figure 4.

**Fig. 4:**
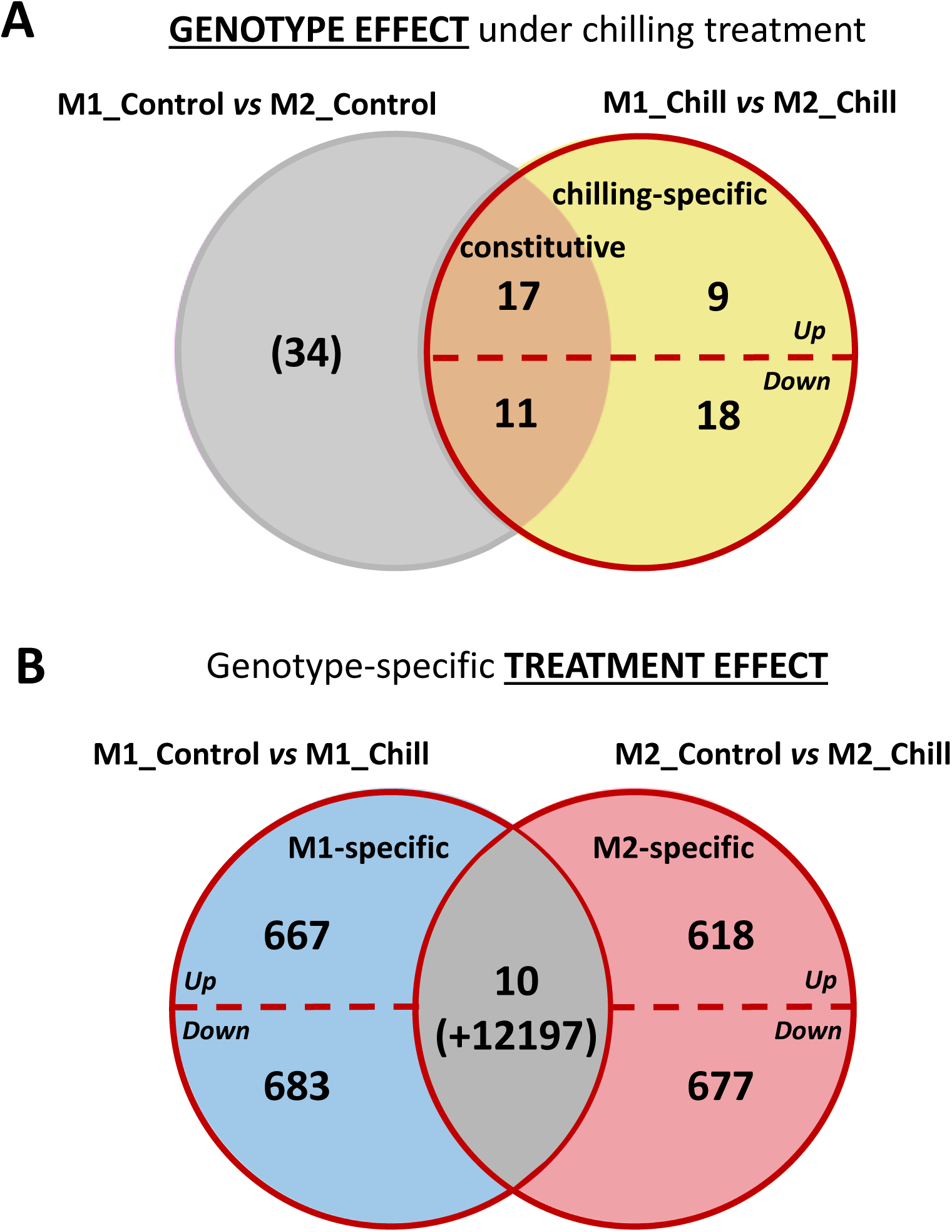
Venn diagrams with an overview of the number of genes whose expression were significantly modulated according to A) the genotype effect and B) the chill-treatment effect. *The analyzes were performed on maize leaves of M1 chill-resistant and M2 chill-sensitive genotypes subjected either to control or chilling treatment*.

Under chilling conditions, 55 genes were differentially expressed between M1 and M2 (Figure 4A; Table S8). Approximately half of these genes were specifically differentially expressed under chilling, whereas the remaining genes were also differentially expressed under control conditions and were therefore considered constitutively differentially expressed. Notably, 27 of the 55 differentially expressed genes (DEGs) were located within the 5 Mb introgressed region (Table S8), and 14 were located within the LD_COL4 region identified by GWAS (Table 4). Among these, only one gene, *Zm00001d048612*, contained SNPs significantly associated with early vigor. This gene was located 1.6 kb upstream the most significant GWAS peak as indicated above. Quantitative RT-PCR analysis confirmed RNA-seq results for 67% of the DEGs located in the introgressed region, including *Zm00001d048612* (Table S9).

**Table 4.** Differentially expressed genes (DEGs) linked to the genotype-effect under chilling treatment that are located in the divergence region in chromosome 4. Asterics (*) shows genes that have been validated by RT-qPCR. The genes whose response is constitutive under both treatments have a symbol (+) or (-) depending on whether they were induced or repressed under the control treatment between the genotypes.

| Gene ID (v4.39) | Chromosomal location (start) | Chromosomal location (end) | Description | LogFC (M1Cold/M2Cold) | Includes GWAS SNP |
| --- | --- | --- | --- | --- | --- |
| Zm000001d048639 | 2 460 060 | 2 463 705 | Disease resistance protein RPM1* | 12.474(+) | No |
| Zm000001d048637 | 2 421 428 | 2 434 362 | Disease resistance RPP13-like protein 4* | 11.774(+) | No |
| Zm000001d048686 | 3 251 054 | 3 252 592 | Uncharacterized protein * | 10.330(+) | No |
| Zm000001d048722 | 4 187 981 | 4 189 310 | Protein SHOOT GRAVITROPISM 5 | 10.011(+) | No |
| Zm000001d048613 | 1 294 081 | 1 306 354 | Disease resistance protein RGA2* | 9.428(+) | No |
| Zm000001d048698 | 3 500 822 | 3 505 327 | Uncharacterized protein * | 7.355(+) | No |
| Zm000001d048635 | 1 863 484 | 1 867 128 | Disease resistance protein RPM1* | 6.566(+) | No |
| Zm000001d048684 | 3 237 295 | 3 249 838 | Cysteine-rich receptor-like protein kinase 8* | 2.080(+) | No |
| Zm000001d048687 | 3 254 797 | 3 257 381 | Heat shock 70 kDa protein 3* | 1.993 | No |
| Zm000001d048592 | 919 275 | 920 997 | RUBISCO activase* | 1.853(+) | No |
| Zm000001d048741 | 4 796 506 | 4 825 085 | Nuclear pore complex protein NUP107 | 1.627(+) | No |
| Zm000001d048606 | 1 119 071 | 1 120 989 | RHOMBOID-like protein* | 1.481(+) | No |
| Zm000001d048696 | 3 440 983 | 3 444 303 | Pectin lyase-like superfamily protein* | 1.343(+) | No |
| Zm000001d048634 | 1 846 584 | 1 848 071 | DIBOA-glucoside dioxygenase BX6 | 1.248 | No |
| Zm000001d048623 | 1 532 821 | 1 534 194 | MYB transcription factor* | 1.018 | No |
| Zm000001d048721 | 4 184 199 | 4 187 807 | Lipase* | -1.283(-) | No |
| Zm000001d048612 | 1 286 634 | 1 292 972 | Vesicle-associated protein 1-1* | -1.404(-) | Yes |
| Zm000001d048615 | 1 368 531 | 1 369 329 | Putative membrane lipoprotein | -1.462(-) | No |
| Zm000001d026713 | 1 528 937 | 1 529 212 | Uncharacterized protein | -1.562 | No |
| Zm000001d048669 | 3 057 593 | 3 060 877 | Auxin-induced beta-glucosidase | -1.586 | No |
| Zm000001d048705 | 3 672 326 | 3 674 403 | Benzoxazinone synthetase* | -1.818 | No |
| Zm000001d048737 | 4 611 294 | 4 613 053 | Uncharacterized protein | -2.257 | No |
| Zm000001d048614 | 1 307 130 | 1 309 817 | Putative GTP-binding protein OBG mitochondrial* | -2.295(-) | No |
| Zm000001d048644 | 2 546 674 | 2 549 655 | NEP-interacting protein 1* | -2.400(-) | No |
| Zm000001d048625 | 1 562 954 | 1 564 019 | O-methyltransferase ZRP4* | -3.141(-) | No |
| Zm000001d048709 | 3 833 329 | 3 835 353 | Indole-3-glycerol phosphate lyase, chloroplastic | -4.335(-) | No |
| Zm000001d048747 | 4 878 850 | 4 884 717 | Uncharacterized protein | -8.709(-) | No |

Treatment effects were then compared between genotypes (Figure 4B). More than 12,000 genes were differentially expressed between chilling and control conditions in both genotypes. In addition, 2,645 genes were differentially expressed in only one genotype, and ten genes were differentially expressed in opposite directions between genotypes. These responses were considered genotype-specific. The corresponding 2,655 genes were distributed across all chromosomes, with approximately 10% located on chromosome 4 (Table S10). Interestingly, none of these genotype-specific DEGs was located within the LD-COL4.1 region, while several were located on chromosome including the LD_COL4.2 region (Figure 5).

**Fig5:**
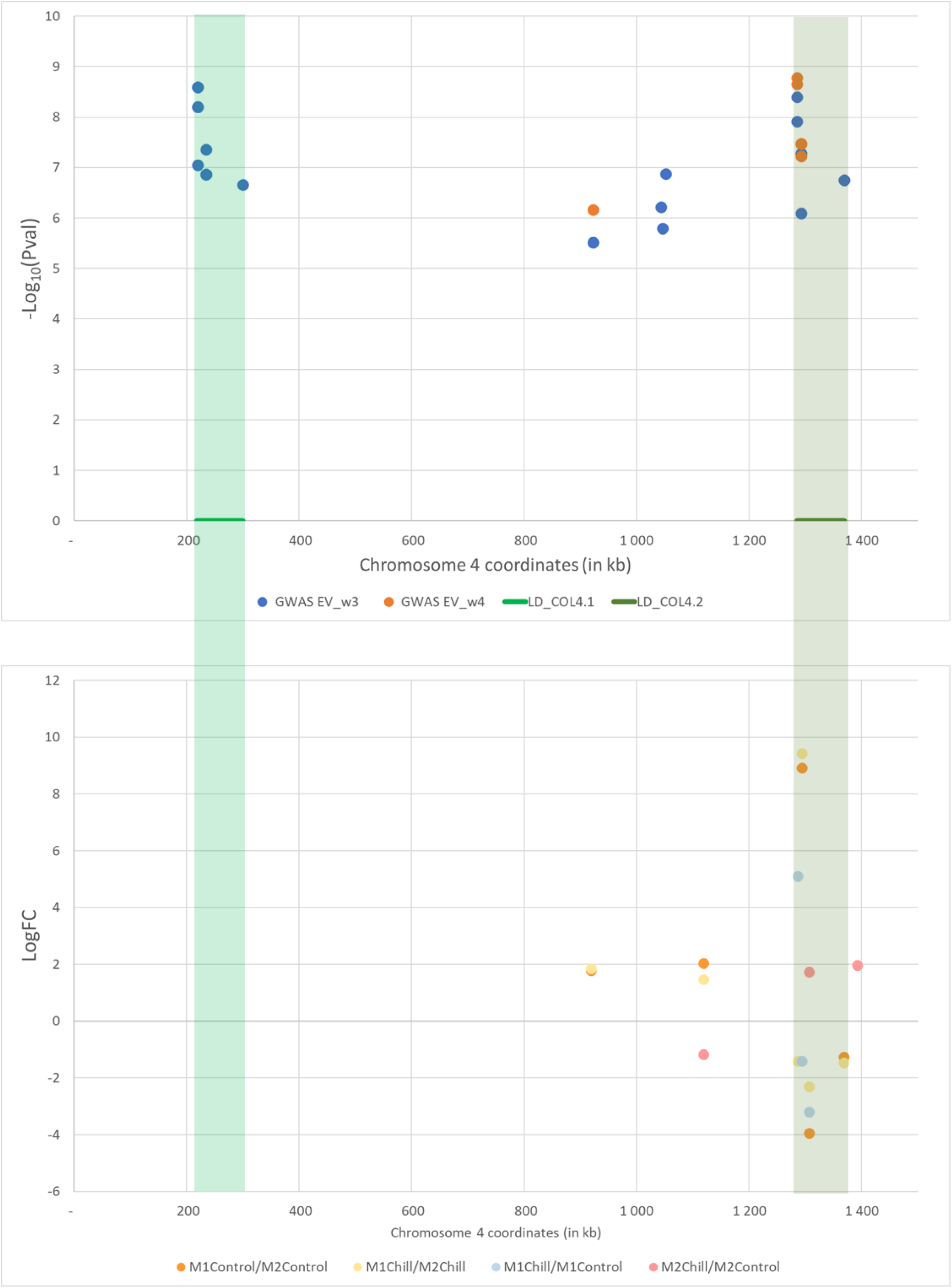
Zoom on chromosomal position of LD_COL4-related SNPs (A) and DEGs (B)

**Fig.6:**
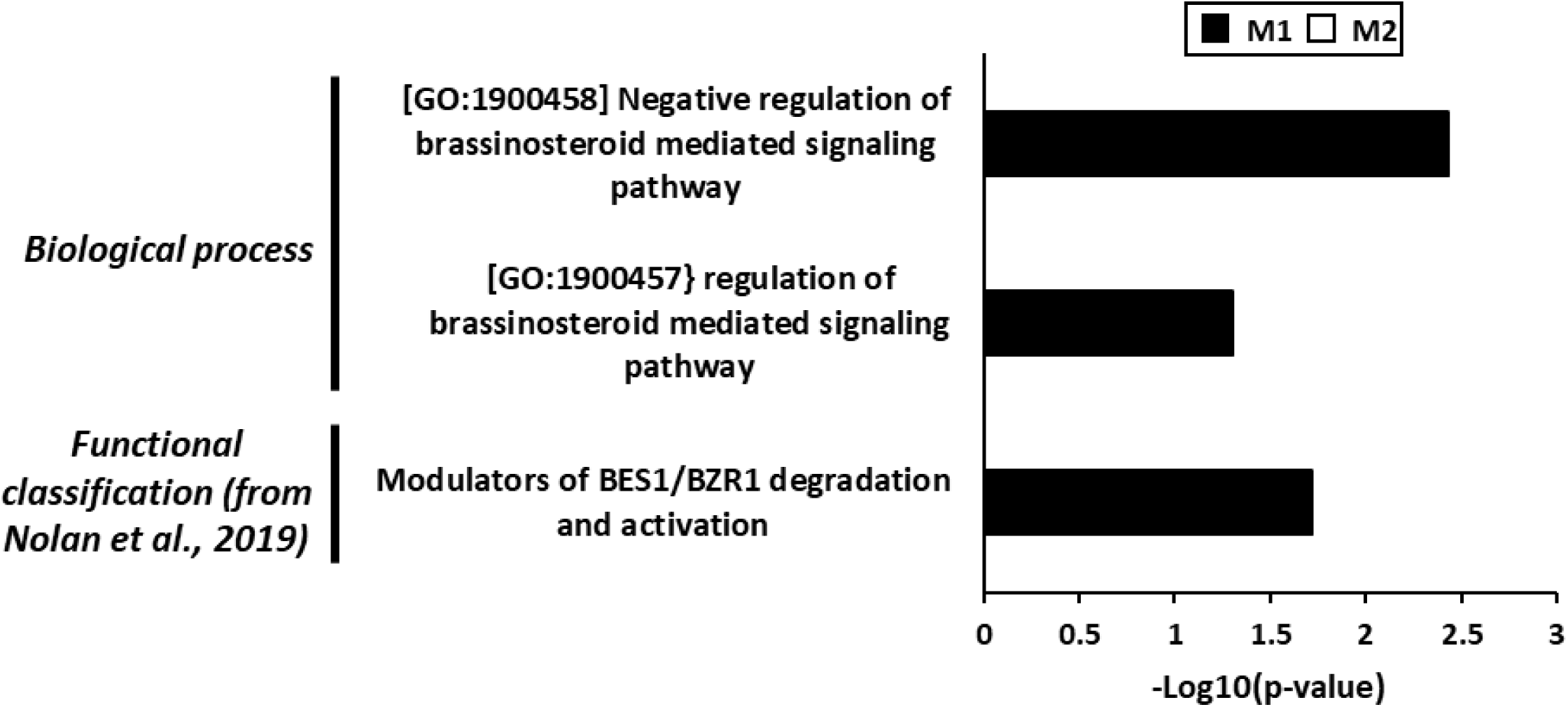
Histogram of enriched biological processes and functional classification related to brassinosteroid of differentially expressed genes (DEGs) regarding the genotype-specific treatment effect*. The functional classification is based on Nolan et al., 2019. The black and white bars correspond to the M1 and M2 genotype, respectively*.

### Brassinosteroid signaling emerges as a contributor to chilling tolerance

The two most significant GWAS peaks were associated with *Zm00001d048582* and *Zm00001d048612* in the LD_COL4.1 and LD_COL4.2 regions, respectively. The Arabidopsis ortholog of *Zm00001d048582,* AT3G09070, also known as OCTOPUS, encodes a polarly localized membrane-associated protein that regulates phloem differentiation (TAIR) and is a positive regulator of the brassinosteroid signaling pathway (Anne et al., 2015). Although this gene was filtered out of the differential expression analysis due to low expression levels, two reads were detected in M1 under chilling conditions. The second gene, *Zm00001d048612,* encodes a serine/threonine protein kinase (Vesicle-associated protein 1-1) and is considered as a brassinosteroid-signaling kinase (BSK) as its rice ortholog, Os04g58750, also known as OsBSK3, promotes BR signaling (Zhang et al., 2016; Kim et al., 2025). This gene was downregulated in M1 compared to M2 under chilling conditions (Table 4), and was also specifically upregulated in M1 under chilling relative to control conditions (Table 5).

**Table 5.**
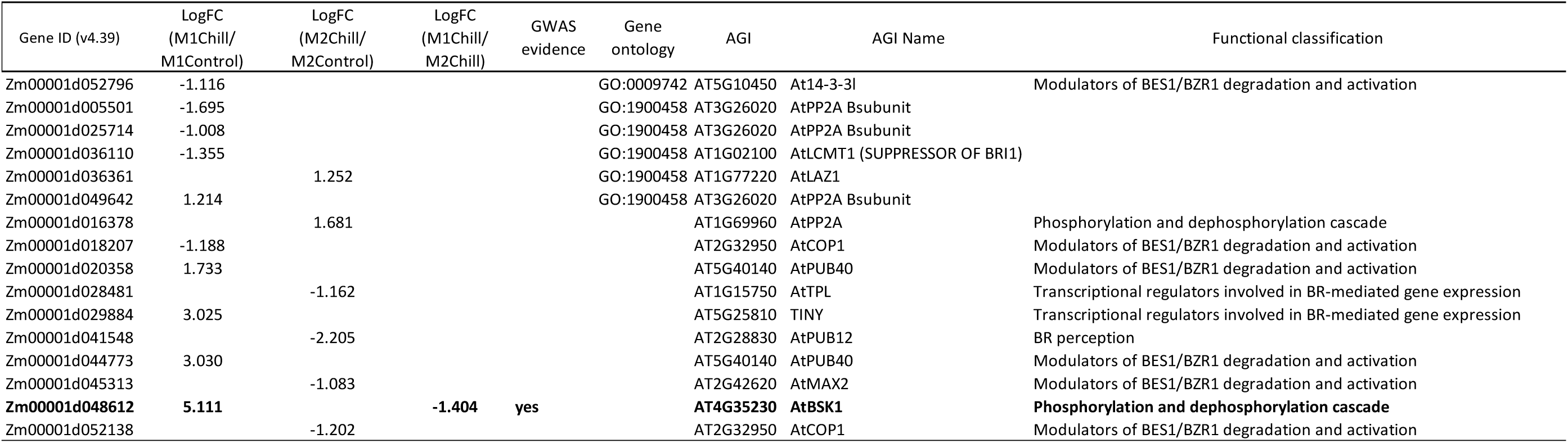
Differentially expressed genes (DEGs) linked to the genotype-specific treatment-effect that are related to the Brassinosteroids. The gene ontologies are from iortho. The functional classification is based on Nolan et al., 2020.

Given that the strongest positional and expression candidates were related to BR signaling pathway, we investigated whether additional genes involved in BR signaling-related were present among the DEGs. Maize genes annotated with BR signaling-related Gene Ontology (GO) terms are listed in Table S11, and maize orthologs of Arabidopsis BR signaling pathway described by Nolan et al. (2020) are provided in Table S12. Among these, sixteen genes were identified among the genotype-specific, chilling-responsive DEGs (Table 5). Gene ontology enrichment analysis revealed that the term GO:1900458 (negative regulation of brassinosteroid mediated signaling pathway) was significantly enriched among the M1-specific DEGs. Three genes (*Zm00001d005501, Zm00001d025714, Zm00001d036110*) were down-regulated and one (*Zm00001d049642*) was up-regulated in M1 under chilling conditions compared with control conditions. In addition, genes classified as modulators of BES1/BZR1 degradation and activation were well represented, with two down-regulated genes (*Zm00001d018207, Zm00001d052796*) and two up-regulated genes (*Zm00001d020358, Zm00001d044773*) in M1. Two genes from this functional class were also specifically down-regulated in M2 (*Zm00001d045313, Zm00001d052138*).

## Discussion

To our knowledge, this study represents the first maize GWAS targeting tolerance to long-term chilling treatment from emergence through the establishment of fully acquired autotrophic growth, although genetic variation for this trait has long been recognized (Hardacre and Eagles, 1980). Previous GWAS focused on germination (Ma et al., 2023; Zhang et al., 2021; Hu et al., 2017; Huang et al., 2013) or very early stages (Yi et al., 2021; Ti et al., 2020; Revilla et al., 2016; Strigens et al., 2013) did not identify our main target region on chromosome 4. Chilling tolerance is known as a complex trait based on multiple independent mechanisms (Sowiński et al., 2025). Responses to chilling stress depends on the developmental stage and stress duration (Burnett 2022) so that genetic determinants of these responses might also be different. This result would partly explain the weak correlation generally observed between performance in growth chamber and field experiments (Jompuk et al., 2005; Strigens et al., 2013) as single-stage growth chamber assays are unlikely to reproduce the complex, dynamic temperature regimes of field conditions. Accordingly, breeding for enhanced chilling tolerance must consider crop phenology and target environment (Burnett et al., 2022). Moreover, identification of LD_COL4 region relied on a few tens inbred lines carrying the alleles conferring a higher chilling susceptibility. In panels with less such inbred lines, the region would likely remain undetected.

LD_COL4 region appeared to carry interesting polymorphisms related to chilling tolerance. While none of the DEG was found below LD_COL4.1 in our dataset, phenotypic effects might be due either to gene expression variation at earlier developmental stages or to deleterious mutations/post-transcriptional changes. *Zm00001d048582* is known to be highly expressed in young plantlets or tissues (SAM, immature tassel and cob) compared to mature leaves (Hoopes et al., 2019), explaining its low read counts in our dataset. However, as the best significant SNPs were found within the coding sequence of *Zm00001d048582*, protein sequence differences might be expected between B73 types and CM174 or Mo17 types. Deleterious mutations or alternative splicing have been shown to contribute to environmental adaptation (Dwivedi et al., 2023; Alhabsi et al., 2025). For example, natural loss of function mutations of the vernalization gene *VRN2* in wheat has led to spring forms (Yan et al., 2004). Similarly, Arabidopsis loss-of-function mutants of OPS, the ortholog of *Zm00001d048582*, are impaired in phloem differentiation and show altered vascular network in cotyledons and intermittent phloem differentiation in roots (Anne et al., 2015). OPS positively regulates BR signaling pathway upstream of the key transcription factors BES1 and BZR1 by confining BIN2 to the plasma membrane and blocking its interaction with BES1/BZR1, thereby influencing flavonoid production and switching from growth to photoprotection under UV-B stress (Liang et al., 2020), an interesting property that could help for chilling tolerance as well. While we could not directly demonstrate Zm00001d048582’s role upstream of BES1/BZR1, six orthologs of BES1/BZR1 modulators were differentially expressed between genotypes in response to chilling, supporting this hypothesis.

*Zm00001d048612*, the BSK underlying LD_COL4.2, was differentially expressed as the three other genes of that region. The phenotypic effect of that region was probably due to a higher transcriptional activity, as the most significant SNPs were found 1.6 kb upstream of the 4 DEG present in the region. In maize lines contrasted for chilling tolerance, chromatin accessibility differences and potentially expressed genes and cis-regulatory sequences open for interaction with transcription factors were identified (Jonczyk et al., 2020), suggesting that LD_COL4.2 could correspond to such a region. In maize, ∼40% of phenotypic variability for agronomically important traits is accounted for by genetic variation within cis-regulatory region (Marand et al., 2025). An eQTL involved in *Zm00001d048612* transcript levels in a panel of 224 maize accessions was found nearby the structural gene, with its effect modulated by drought stress (Liu et al., 2020). In Arabidopsis, BSKs are phosphorylated and activate BSU1 phosphatase to inhibit BIN2, demonstrating conserved BR signaling mechanisms (Nolan et al., 2020).

BR signaling appears to be involved in chilling tolerance in several species like tomato (Wang et al., 2022; Fang et al., 2021), sorghum (Mu et al., 2022) and rice (Fang et al., 2019). These observations reinforce our hypothesis of the role of *Zm00001d048582* and *Zm00001d048612* in chilling tolerance in maize. However, none of these two genes has been reported in previous chilling tolerance experiment (listed by Ma et al., 2022), probably because of the specificity of the CM174 allele and the type of chilling stress imposed. Meng *et al* (2021) show that gene expression responses to cold stress vary even among genotypes within the same species. This regulation relies on coordinated interactions between cis- and trans-regulatory elements, transposable elements, and epigenetic mechanisms. Considering the potential role of both *Zm00001d048582* and *Zm00001d048612* candidates in the same signaling pathway, it can be expected epistatic interactions between both loci, making LD_COL4 a valuable target for dissecting such interactions.

In maize, BR signaling is also known to be involved in tolerance to other abiotic stress like drought (Castorina et al., 2018). BR homeostasis and signaling is considered as a way of improving the adaptation to environmental stress (Zolkiewicz and Gruszka, 2025). But it is also involved in other agronomic traits like plant architecture (Zhang et al., 2025). Among plant architectural traits, plant height dynamic might be of considerable interest in the case of agroecology practices like intercropping or pesticides reduction, as an element of competition. Plant height and more specifically dwarfism phenotypes are traits that can be mediated through BR signaling. It is the case for example in maize with *GRAS42* (Kaur et al., 2024), in sorghum with *dw1* (Hirano et al., 2017), in rice with *ebisu dwarf* (Hong et al., 2003), in barley with *HvDWARF* (Sadura et al., 2019). In sorghum, Marla et al (2019) found that a QTL for chilling tolerance after early planting in the field mapped close to *dw1* and *dw3*, two genes massively used in grain sorghum during the green revolution to shorten plants. Selection for dwarfism may have counter-selected linked chilling-tolerance alleles, although pleiotropic effects of these loci cannot be excluded. Plant height plasticity in response to diurnal temperature range is strongly influenced by genotype × environment interactions, with consistent reaction norm QTLs identified at *Dw1*, *Dw3*, and *qHT7.1* (Mu et al., 2021). *Dw1* encodes a BR signaling component (Hirano et al., 2017), while *Dw3* and *qHT7.1* are orthologous to maize *Br2* (auxin transporter) and *Br1* (MYB transcription factor) respectively (Jiao et al., 2023). The interconnection between BR signaling pathway and plant height plasticity provides new opportunities to breeder and geneticist to tackle interesting adaptation alleles to field environments, taking advantage of high throughput phenotyping tools using unmanned aerial vehicle.

In conclusion, our study identifies LD_COL4 as a key genomic region for long-term chilling tolerance in maize, with *Zm00001d048582* and *Zm00001d048612* emerging as promising candidate genes potentially acting through BR signaling. These findings highlight the importance of developmental stage-specific responses and the complex genetic architecture underlying chilling tolerance. The result provides valuable targets for breeding programs aiming to improve early vigor under early-planting and offer a foundation for functional studies to dissect the molecular mechanisms and potential epistatic interactions between loci.

## Acknowledgements

This research was jointly supported by the French National Research Agency (ANR), the German Federal Ministry of Education and Research (BMBF), and the Spanish ministry of Science and Innovation (MICINN) as the CORNFED project, by the French National Research Agency (ANR) and France Agrimer as the AMAIZING (ANR-10-BTBR-01) project. We are especially grateful to Alain Charcosset, Laurence Moreau, Peter Rogowsky for the management of AMAIZING project and work packages, to Jean-François Hû, Dominique Rabier for their help with maize culture, sampling, and phenotyping, and to Cyril Bauland, Jacques Laborde, Stéphanie Castel for their contribution to plant material building. We also thank Christine Coffigniez and Martine Thomas for their support in the daily administrative work.

## Supporting information

**FigS1:**
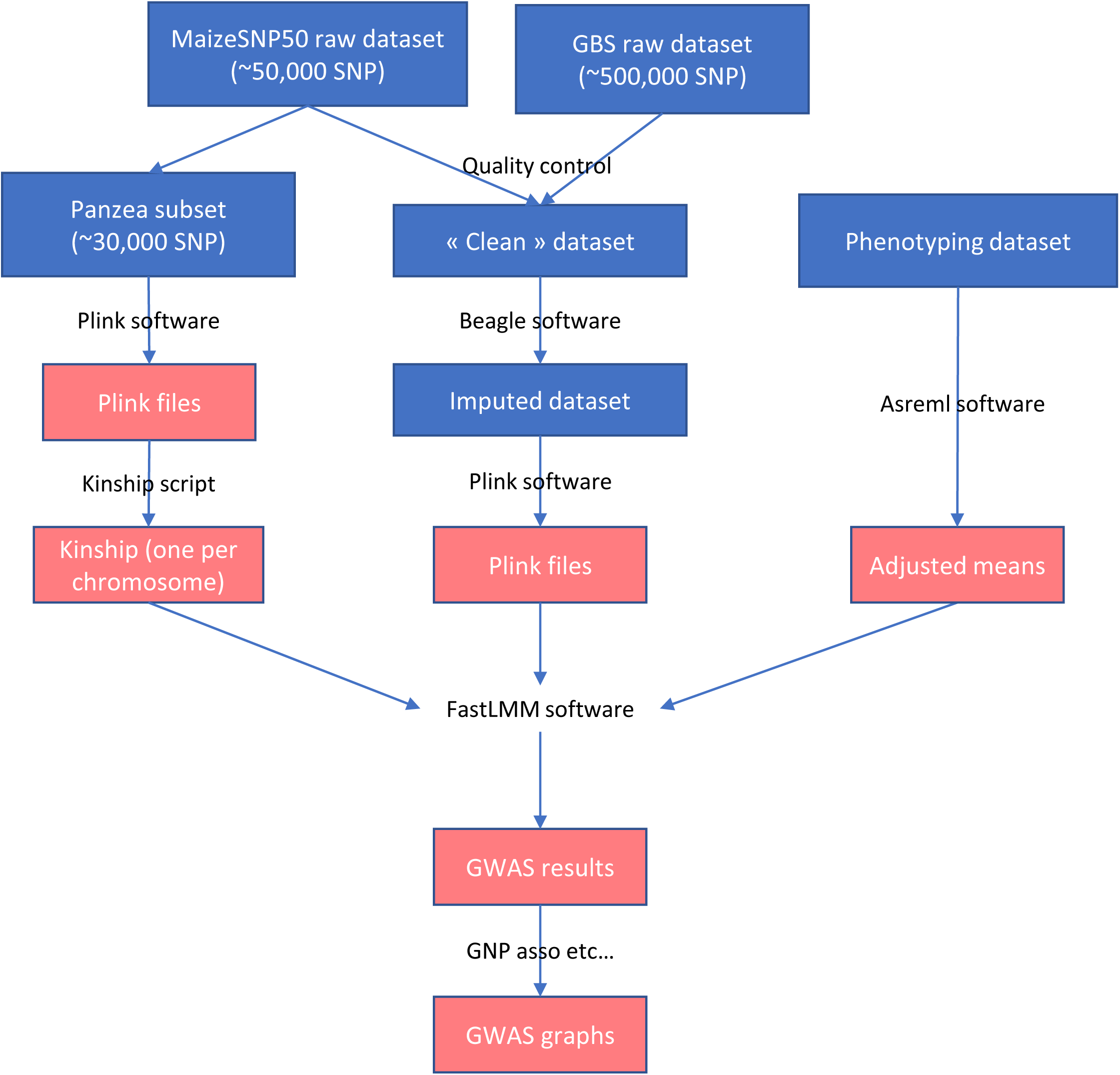
The analysis pipeline of the genome-wide association study.

**Fig.S2:**
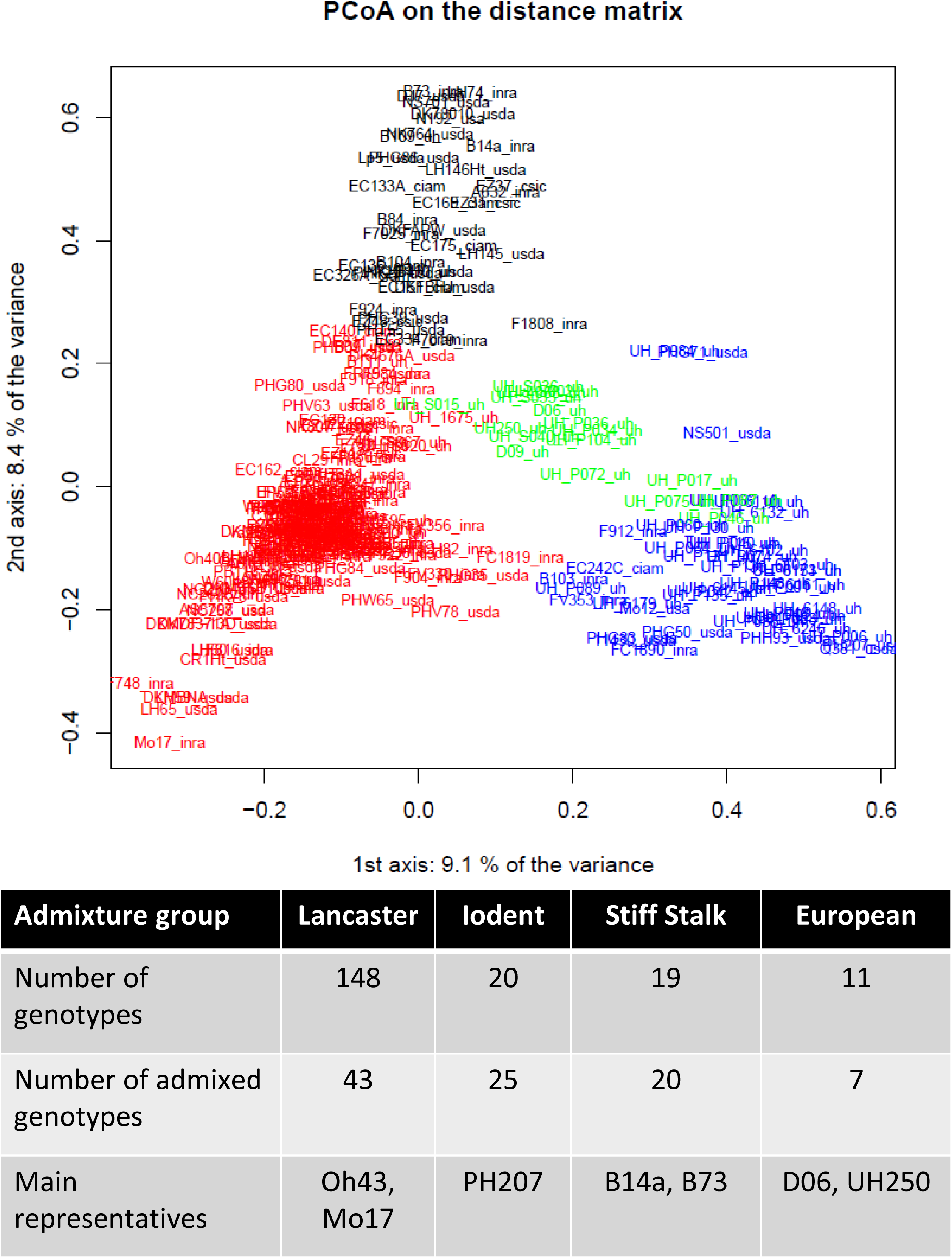
Admixture groups on first PCA components based on Panzea SNPs et number of representatives in the diversity panel.

**FigS3:**
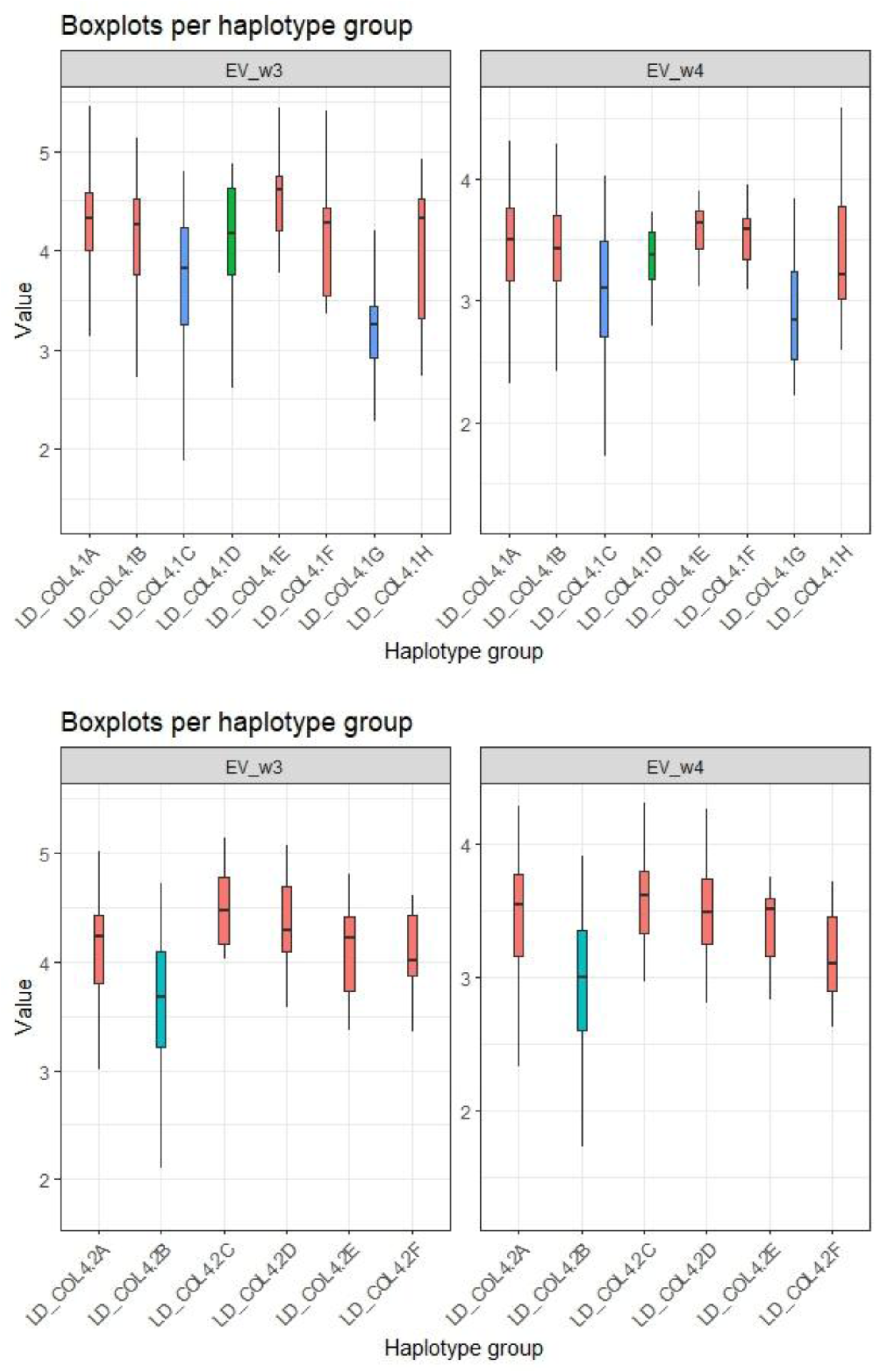
Main haplotype effects on early vigor evaluated after three (w3) or (faur (w4) weeks of chilling treatment.

Table S2: Best Linear Unbiased Estimates (BLUEs) for all traits and genotypes

Table S3. Polymorphisms significantly associated with phenotypic traits under chilling conditions: positions, GWAS results and related gene models from Zea_mays.B73_RefGen_genes.v4_39.

Table S4: Genotyping and haplotypes for SNP delineating the two genomic regions, LD_COL4.1 and LD_COL4.2, associated with earlyvigor under chilling temperatures.

Table S5: Flanking sequences of SNPs used for QTL-NILs construction.

Table S6. Total count of RNA sequencing of leaf samples of two QTL-NILs grown under chilling (CHILL, 16°C day/14°C night) and control (CONTROL, 25°C day/ 22°C night) conditions in growh chamber. M1 is the chilling tolerant NIL, similar to B73 inbred line, while M2 is the chilling sensitive NIL, carrying a CM174 segment of approx 5 Mb on chromosome 4.

Table S7. SNP-associated candidate genes for early vigor and leaf width under chilling conditions and mapping results in RNAseq experiment on QTL-NILs.

Table S8. Differentially expressed genes (DEGs) linked to the genotype-effect under chilling.

Table S9. Primer to validate differentially expressed genes (DEGs) linked to the genotype-effect in M1 and M2 genotype and located in the divergence region of chromosome 4 using RT-qPCR. In red, are the primers for the housekeeping genes.

Table S10: Differentially expressed genes (DEGs) linked to the treatment-effect specifically for M1 genotype or M2 genotype.

Table S11: Maize genes with GO terms associated with brassinosteroid signaling pathway.

Table S12: Maize orthologs of Arabidopsis genes involved in BR signaling pathway (list from Nolan et al., 2020)

